# Dysfunctional parvalbumin interneurons in a genetic mouse model of schizophrenia

**DOI:** 10.1101/2023.09.10.557034

**Authors:** S. Hijazi, M. Pascual-García, A. Tolido, A. Pham, S.A. Kushner

## Abstract

The 22q11 deletion syndrome (22q11DS) is an interstitial microdeletion associated to an increased risk of developing schizophrenia. In this disorder, there is a dysfunction in the overall connectivity of the brain. Parvalbumin-expressing (PV^+^) interneurons have been associated with multiple pre- and post-synaptic impairments that affect various brain regions. Specifically, previous results have suggested that alterations in hippocampal networks may be related to PV^+^ interneurons dysfunction. In this study, we used the Df1 mouse model that carries the 22q11 deletion to examine the excitability of PV^+^ cells in the dorsal CA1 region of the hippocampus, due to its importance in memory and cognition. We found that PV^+^ interneurons were hyperexcitable in this region. To understand the source of the altered excitability, we measured potassium currents, highly involved in the intrinsic firing properties of neurons. We observed that voltage-gated potassium channel subfamily A member 1 (K_v_1.1) was impaired in PV^+^ cells. Specific activation of this channel recovered some of the excitability disturbances observed in Df1 mice. Furthermore, blockade of synaptic inputs also restored PV^+^ interneuron’s excitability. Taken together, these results suggest that PV excitability is increased in the CA1 region of the hippocampus and it is partially mediated by K_v_1.1 in a mouse model of 22q11DS.

## Introduction

The 22q11 deletion syndrome (22q11DS) is a neurodevelopmental disorder in which neuronal circuitry and behavioural paradigms are disrupted. The size of the interstitial deletion ranges between 1.5 to 3 Mb^1^, and no correlation between the size of the deletion and the phenotype has been found^1^. This microdeletion is widely associated with an increased risk of schizophrenia. 25% of the carriers develop the disease^2^, which accounts for 1-2% of the total cases with schizophrenia^3,4^. Approximately 60 genes are located in the 3 Mb microdeletion area^5^, although ∼30 genes found in the 1.5 Mb seem to be critical for the development of the disease^6^. Most of these genes are located, but not exclusively, in the brain^7^. Genes in this region have been individually studied, and animal models of the 22q11DS have been developed to understand the role of the cluster on neuronal maturation and circuit functioning.

Schizophrenia is a polygenic disease characterised by the dysfunction of brain networks^8^. Many areas present functional and morphological changes, but the hippocampal-prefrontal pathway is particularly affected^9,10^. Communication between the hippocampus and the prefrontal cortex (PFC) is crucial for working memory, cognition, executive function and emotions^9^, which are dysregulated in schizophrenia^11,12^. Altered neurodevelopmental processes provide a robust framework for the aetiology and early onset of the carriers of the 22q11DS mutation^6^. However, it is still unknown which are the fundamental mechanisms that underlie the pathophysiology of 22q11DS and schizophrenia.

Fast-spiking parvalbumin (PV^+^) interneurons are GABAergic cells that play a key role in the regulation of network dynamics and synchronisation^13,14^. Research has pointed out its causal role in schizophrenia^15–17^. Developmental impairments in PV^+^ interneurons in 22q11DS have been shown in different cortical and hippocampal areas^18,19^. Hyperactivity of hippocampal structures in schizophrenia^20,21^, converging with deficits in PV^+^ number and activity^22–25^, has been linked to the onset and progression of the disease^24–26^. Nevertheless, the link between the microdeletion and PV^+^ interneuron functionality is poorly understood.

PV^+^ interneuron plasticity is largely regulated, among others, by synaptic inputs and intrinsic mechanisms dependent on specific potassium channels^27^. K_v_1 and K_v_3 channels are important regulators of neuronal excitability^27–30^. Loss of function mutations or transcriptional changes in these channels may have an effect on the near-threshold excitability and the repolarisation of the action potential, as well as for the firing rates and synaptic transmission of PV^+^ interneurons^29,31,32^. Notably, both of the genes encoding for these potassium voltage-gated channels are altered in the transcriptome of schizophrenia patients^33^. The alteration in gene expression might, in turn, provoke a dysregulation of the PV^+^ intrinsic properties and presynaptic activity and therefore of the local network excitability.

Here, we studied the electrophysiological properties of PV^+^ interneurons using a genetic mouse model containing the orthologous 22q11DS, named Df1. Strikingly, we found that PV^+^ interneurons in the CA1 region of the hippocampus, but not in the PFC, are hyperexcitable. When synaptic activity was blocked, PV neurons excitability was restored, suggesting that synaptic activity plays a role in the observed hyperexcitability of PV neurons in Df1 mice. To deepen our understanding on the mechanisms underlying this hyperexcitability, we studied voltage-gated potassium channel subfamily A member 1 (K_v_1.1), present in the axon initial segment and the presynaptic sites of these cells. We observed that these channels were hypofunctional, suggesting an intrinsic mechanism that drives the perturbations in excitability. Interestingly, activation of K_v_1.1 using a specific agonist, brought electrophysiological parameters to baseline levels. Lastly, we studied the myelination of PV^+^ interneurons, important for their proper functioning, and concluded that it was not affected in the Df1 model.

## Results

### Hippocampal PV^+^ interneurons show early increased excitability in young adult Df1 mice

To selectively investigate the excitability of PV^+^ interneurons in 22q11DS young-adult mice, we crossed Df1 mice with transgenic PV::Ai14 mice. Next, we performed patch-clamp recordings and biocytin-filling of the cells. Whole-cell recordings of PV^+^ interneurons in PFC did not show any sign of abnormal excitability (**Supplementary figure 1**). Conversely, in dorsal CA1 (dCA1) region of the hippocampus, PV^+^ interneurons displayed an increased excitability represented by a reduced action potential (AP) threshold (**Figure 1c**), afterhyperpolarisation amplitude (AHP; **Figure 1g**), and rheobase (**Figure 1e**), while their membrane potential (RMP) was unaltered (**Figure 1a**). The rest of the active and passive properties were unaffected (**Figure 1a-h**). Firing frequency of PV^+^ interneurons, however, was increased at lower depolarisation steps (**Figure 1i, j**), indicating that hippocampal PV^+^ interneurons in Df1 are hyperexcitable.

**Figure 1.**
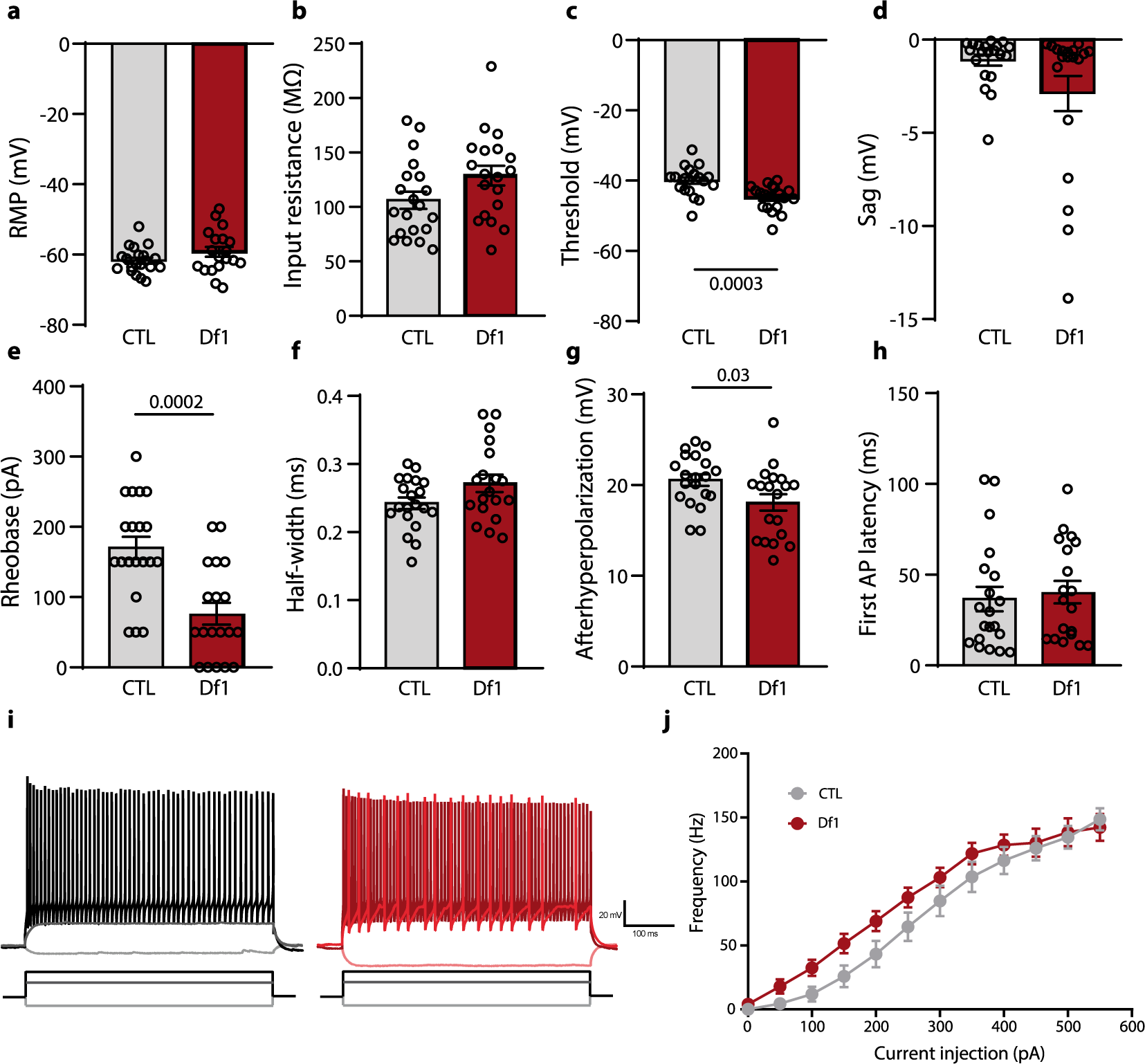
Increased excitability of hippocampal PV^+^ interneurons in Df1 mice. **a)** Resting membrane potential was not affected in Df1 PV^+^ interneurons (RMP; Mann-Whitney: U = 142; p-value = 0.18; CTL average: -61.71 mV; Df1 average: -59.24). **b)** Input resistance was not changed in Df1 PV^+^ interneurons (t-test: t = 1.87; p-value = 0.069; CTL average: 106.2 MΩ; Df1 average: 128.7 MΩ). **c)** AP threshold is reduced in Df1 PV^+^ interneurons (t-test: t = 3.96; p-value = 0.0003; CTL average: -40 mV; Df1 average: - 44.89 mV). **d)** Sag is not affected in Df1 PV^+^ interneurons (Mann-Whitney: U = 138; p-value = 0.15; CTL average: -1.1 mV; Df1 average: -2.9 mV). **e)** Rheobase is decreased in Df1 PV^+^ interneurons (t-test: t = 4.2; p-value = 0.0002; CTL average: 170 pA; Df1 average: 76.32 pA). **f)** Half-width is not significantly affected in Df1 PV^+^ interneurons (t-test: t = 1.91; p-value = 0.063; CTL average: 0.24 ms; Df1 average: 0.27 ms). **g)** The afterhyperpolarisation (AHP) is reduced in Df1 PV^+^ interneurons (t-test: t = 2.25; p-value = 0.03; CTL average: 20.55 mV; Df1 average: 19.1 mV). **h)** The latency of the first elicited AP was not altered in Df1 PV^+^ interneurons (t-test: t = 0.41; p-value = 0.68; CTL average: 36.6 ms; Df1 average: 40.36 ms). **i)** Representative traces of PV^+^ interneurons in control (left) and Df1 (right) cells at -100 pA, 100 pA and 300 pA. n = 20/19 cells, respectively. **j)** Firing frequency in response to depolarising steps from 0 to 550 mV during 500 ms (Two-way ANOVA (group x current); F (11, 407) = 1.83; p = 0.047).

### PV^+^ interneurons excitability is dependent on synaptic activity

To test whether PV^+^ interneuron hyperexcitability in dCA1 was driven by synaptic activity, we exposed hippocampal slices to a cocktail of blockers including CNQX, AP5 and gabazine. Bath application of synaptic blockers resulted in a recovery of PV^+^ interneurons excitability in Df1 mice (**Figure 2a-f**), confirming the involvement of synaptic activity in the excitability of PV^+^ interneurons. In order to explore the effect of the synaptic inputs on neuron excitability, we compared baseline properties of PV^+^ interneurons to their properties when synaptic inputs were blocked. In controls, input resistance increased, in agreement to the expected effect of synaptic blockers. Additionally, rheobase was decreased, caused by the depolarisation of the cells. Contrarily, input resistance and rheobase in mutant cells were unchanged upon blocking synaptic transmission, whilst the AP threshold was decreased, and the shape of the AP varied, highlighting a clear difference in how PV interneurons responded to blocking synaptic activity between DF1 mice and control mice (**Supplementary figure 2**). Surprisingly, we observed no significant alterations in the frequency or amplitude of either spontaneous excitatory or inhibitory postsynaptic currents (sEPSC and sIPSC, respectively) (**Figure 2g-j**).

**Figure 2.**
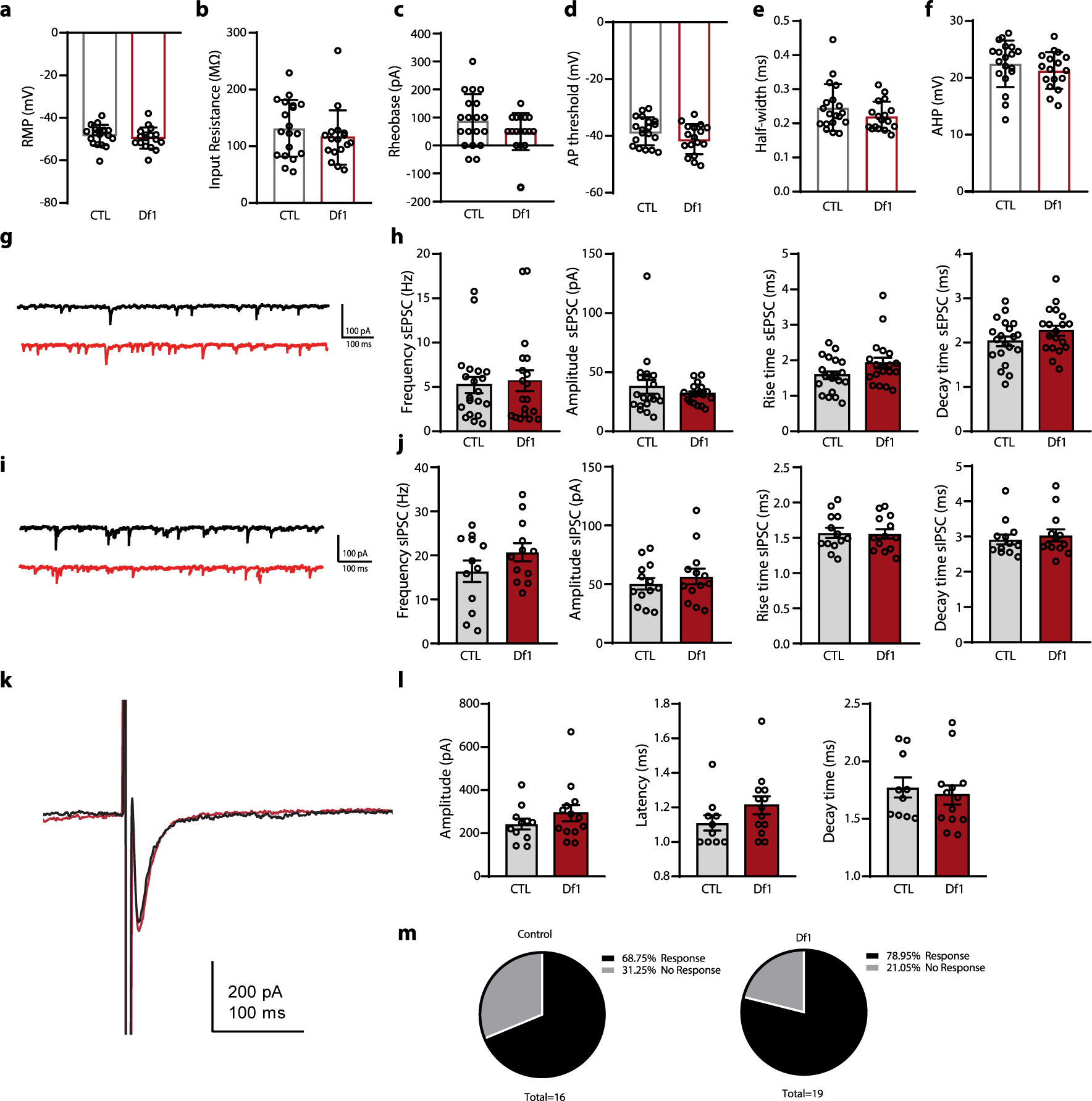
Synaptic properties of PV^+^ interneurons in the hippocampus. **a)** Resting membrane potential was not affected (t-test: t = 0.87; p-value = 0.39; CTL average: -48.11 mV; Df1 average: -49.53). **b)** Input resistance was not changed (t-test: t = 0.99; p-value = 0.33; CTL average: 131.6 MΩ; Df1 average: 115.3 MΩ). **c)** Rheobase is similar between groups (t-test: t = 1.44; p-value = 0.16; CTL average: 89.47 pA; Df1 average: 50 pA). **d)** AP threshold is maintained across groups (t-test: t = 1.7; p-value = 0.11; CTL average: -38.36 mV; Df1 average: -41.12 mV). **e)** Half-width is not changed (t-test: t = 1.3; p-value = 0.204; CTL average: 0.24 ms; Df1 average: 0.22 ms). **f)** AHP is unaltered (t-test: t = 0.96; p-value = 0.34; CTL average: 22.48 mV; Df1 average: 21.29 mV). **g)** Example trace of sEPSC in control (up) and Df1 (down) cells. **h)** The sEPSC properties are not different between Df1 and CTL group (Frequency; t-test: t = 0.32; p-value = 0.75; CTL average: 5.22 Hz; Df1 average: 5.7 Hz; Amplittude; t-test: t = 0.32; p-value = 0.75; CTL average: 37.55 pA; Df1 average: 31.8 pA; rise time; t-test: t = 1.84; p-value = 0.074; CTL average: 1.58 ms; Df1 average: 1.92 ms; decay time; t-test: t = 1.52; p-value = 0.14; CTL average: 2.03 ms; Df1 average: 2.27 ms). **i)** Example trace of sIPSC in control (up) and Df1 (down) cells. **j)** The sIPSC properties are not different between Df1 and CTL group (Frequency; t-test: t = 1.36; p-value = 0.19; CTL average: 16.4 Hz; Df1 average: 20.74 Hz; Amplittude; t-test: t = 0.75; p-value = 0.46; CTL average: 50.3 pA; Df1 average: 56.5 pA; rise time; t-test: t = 0.14; p-value = 0.88; CTL average: 1.57 ms; Df1 average: 1.56 ms; decay time; t-test: t = 1.52; p-value = 0.54; CTL average: 2.92 ms; Df1 average: 3.04 ms). **k)** Example traces of autaptic responses. **l)** Autaptic amplitude did not vary between control and Df1 cells (t-test: t = 1.07; p-value = 0.29; CTL average: 243 pA; Df1 average: 293 pA). Autaptic latency was unaltered across control and Df1 groups (t-test: t = 1.43; p-value = 0.16; CTL average: 1.11 ms; Df1 average: 1.21 ms). Autaptic decay did not differ between cellular group (t-test: t = 0.53; p-value = 0.59; CTL average: 1.77 ms; Df1 average: 1.71 ms). **m)** Percentage of autaptic responses in both groups.

To regulate their own firing, PV^+^ interneurons also synapse onto themselves and this has been shown to be important for the modulation of cortical networks^34^. Autaptic inhibition can be a stronger way of inhibition than any other form of synaptic activity onto PV^+^ interneurons^35^. Hence, we sought to check whether alterations in autaptic strength could be accompanying changes in PV excitability. We found that the percentage of cells with autaptic responses was not affected in the Df1 model (**Figure 2m**). Furthermore, the amplitude, latency and decay of the autaptic transmission were also not altered in Df1 PV^+^ interneurons (**Figure 2k, l**). These data indicate that neither synaptic nor autaptic inputs are affected in PV^+^ interneurons, but synaptic activity is involved in their excitability.

### Hippocampal PV neurons show alterations in Kv1.1 channel expression and function in Df1 mice

We next aimed to elucidate which were the intrinsic mechanisms driving PV hyperexcitability. Voltage-gated potassium channels 1 (K_v_1.1) are important regulators of the high-firing frequency characteristic of PV^+^ interneurons, as well as for the modulation of the action potential threshold^29,36^, both altered in Df1 PV^+^ cells. Moreover, K_v_1.1 channels have been shown to be critically involved in linking intrinsic excitability and synaptic strength^37^. In order to specifically assess the functionality of K_v_1.1 channels in PV^+^ interneurons, we performed voltage-clamp recordings. We determined that potassium currents in PV^+^ interneurons in voltage-clamp were affected (**Figure 3a**). K_v_1.1 channels activate at -40 mV and deactivate at - 10 mV^38^. Using a K_v_1.1 selective blocker dendrotoxin-K (DTx-K), we could isolate K_v_1.1-specific currents in PV^+^ interneurons. We observed a significant reduction in K_v_1.1 current at -30 mV in PV^+^ interneurons from Df1 mice compared to controls (**Figure 3b, c**). Furthermore, at higher voltage steps, potassium currents are higher than in controls, suggesting that other potassium channels activate and are also involved in the potassium current impairments (**Supplementary figure 3**). Together, this suggests that potassium currents are affected, and particularly K_v_1.1 channels are not properly functional.

**Figure 3.**
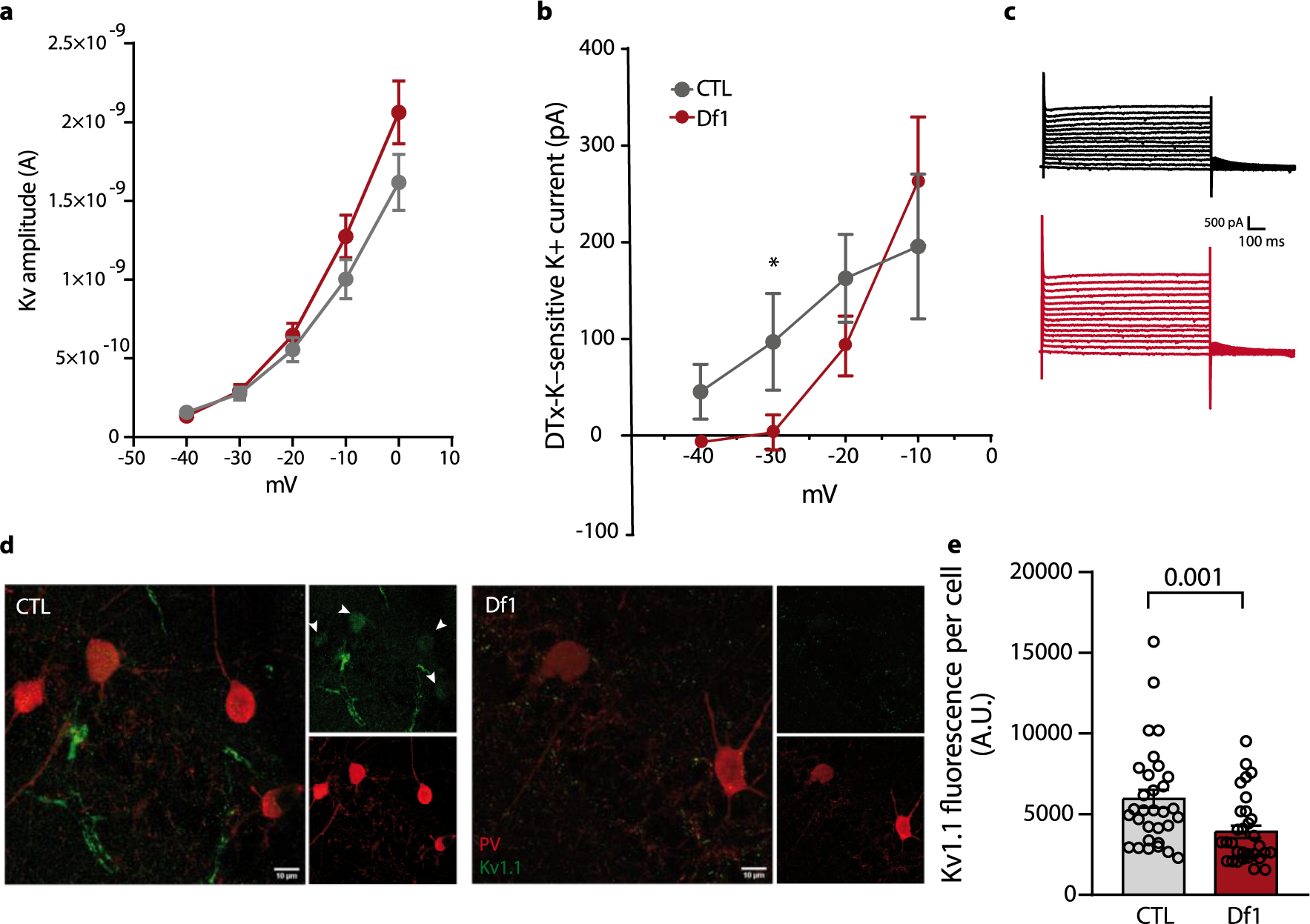
PV^+^ cells show impaired Kv1.1 channel. **a)** Plot of the voltage dependence of the potassium-sensitive currents in Df1 and control PV^+^ interneurons (ANOVA: F (4, 104) = 3.61; p-value = 0.0085; Šidák’s multiple comparisons test; ns). **b)** Plot of the voltage dependence of the DTx-K-sensitive currents in Df1 and control PV^+^ interneurons (ANOVA: F (3, 36) = 2.72; p-value = 0.059; Šidák’s multiple comparisons test; ns; t-test at -30 mV: t = 1.98; p = 0.036). Example traces of voltage DTx-K potassium-sensitive currents. **c)** Example high magnification pictures of colocalization of PV and Kv1.1 in dCA1 in control and Df1 mice. **d)** Kv1.1 fluorescence is significantly reduced in PV cells (Mann-Whitney: U = 252; p-value = 0.0011).

To further estimate whether the expression of K_v_1.1 channels was also altered, we immunostained and quantified the fluorescence of K_v_1.1 channels in PV^+^ interneurons in the dCA1. PV^+^ interneurons had substantially less K_v_1.1 channel fluorescence (**Figure 3d, e**), indicating that these channels are hypofunctioning in PV^+^ interneurons. Western blot analysis of the whole hippocampus showed no alterations in the expression of K_v_1.1, indicating that this alteration was specific to this area (**Supplementary figure 4**). These channels are found both in presynaptic terminals and in the axon initial segment (AIS) of PV^+^ interneurons^29^. The AIS is a complex structure highly enriched with channels to modulate AP generation and hence AP threshold^39^. It is a plastic structure that adjusts its position to adapt to changes in the environment^40^. We measured AIS length to estimate whether structural modification had occurred as a cause or response of changes in PV^+^ interneuron excitability. We observed that the AIS in PV^+^ interneurons had not changed compared to control cells (**Supplementary figure 4**). Therefore, we can conclude that potassium currents are affected in PV^+^ interneurons and K_v_1.1 channel are reduced without changes in the AIS.

### Specific activation of Kv1.1 channels partially restores PV^+^ cells*’* hyperexcitability

With the ultimate aim of restoring PV^+^ interneuron hyperexcitability through K_v_1.1 channel activation, we employed N-[4-(Trifluoromethyl)phenyl]glycine (4-TFMPG), a specific agonist of K_v_1.1 channels^41^. Bath application of the activator increased the AP threshold in the Df1 condition, restoring it to control levels (**Figure 4a**). Intriguingly, no changes in AP threshold were observed in the control group, and in both groups, none of the other intrinsic properties were affected upon activation of K_v_1.1 channels (**Figure 4b-d**). Remarkably, the firing frequency, also modulated by K_v_1.1 channels, was decreased after the application of the activator in Df1 cells, restoring the firing rate to control levels (**Figure 4e, f**). Next, to examine the effect of K_v_1.1 activation when synaptic inputs are blocked, we carried out whole-cell recordings of PV^+^ interneurons with synaptic blockers of AMPA, NMDA and GABA_A_ receptors. Notably, activation of K_v_1.1 channels when synaptic transmission was blocked had no effect io PV^+^ interneurons of Df1 mice but did show an effect in control cells, implicating synaptic K_v_1.1 channels in the observed rescue (**Supplementary figure 3**).

**Figure 4.**
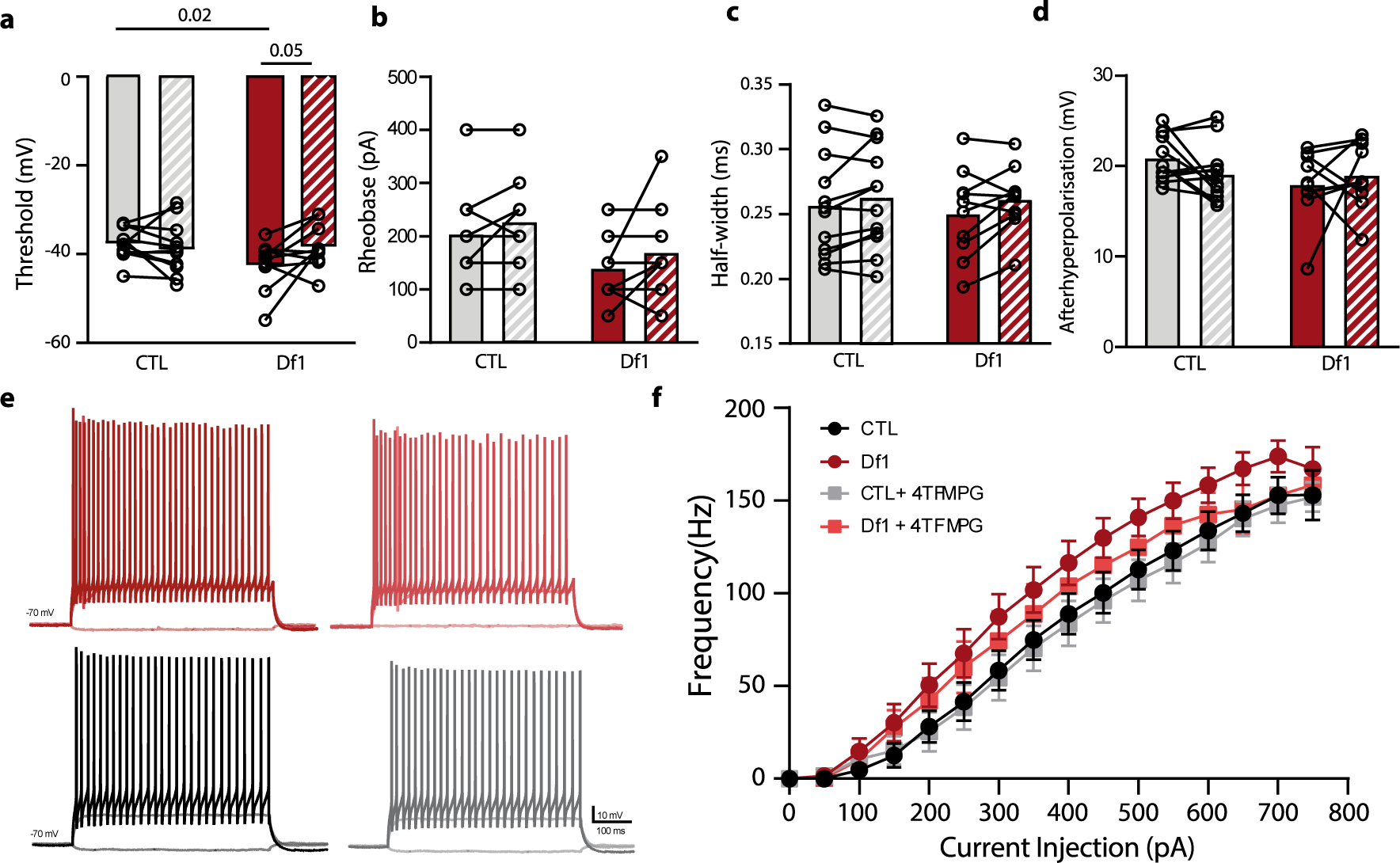
4TFMPG selective agonist for Kv1.1 restores some electrophysiological properties of PV^+^ interneurons. **a)** AP threshold was significantly reduced in the Df1 group after application of 4TFMPG (Two-way ANOVA: F (1,20) = 5.21; p-value = 0.034; Šidák’s multiple comparisons test; CNT, p = 0.72; Df1, p = 0.05). **b)** Rheobase was not significantly increased in either group after application of 4TFMPG (Two-way ANOVA: F (1,20) = 0.08; p-value = 0.77). **c)** Half width was not changed after bath application of 4TFMPG (Two-way ANOVA: F (1,20) = 0.49; p-value = 0.49; Šidák’s multiple comparisons test; CNT, p = 0.34; Df1, p = 0.078) **d)** AHP remained unvaried after application of 4TFMPG (Two-way ANOVA: F (1,20) = 0.14; p-value = 0.49; Šidák’s multiple comparisons test; CNT, p = 0.29; Df1, p = 0.71) **e** Example traces of PV^+^ cells (from left to right) of Df1 baseline, Df1 + 4TFMPG, control baseline and control + 4TFMPG). **f)** Firing frequency in response to depolarising steps from 0 to 750 mV during 500 ms before and after bath application of the 4TFMPG (Two-way ANOVA: F (45, 600) = 1,043).

Taken together, partial recovery of the phenotype was observed upon activation of K_v_1.1, indicating that the hyperexcitability of PV^+^ interneurons in Df1 mice is likely driven by a decrease in K_v_1.1 activity.

### Hippocampal PV^+^ neurons have a normal myelination pattern

Myelination of a cell is highly correlated with its intrinsic excitability^42^. PV^+^ interneurons are myelinated^43,44^, and there is a strong likelihood that their demyelination is an underlying mechanism in schizophrenia^17^. Therefore, using biocytin-filled cells, we examined their myelination profile in Df1 dCA1. A high percentage of PV^+^ interneurons in both control and Df1 mice were myelinated (**Figure 5a, d**) (control: 90%; Df1: 73%). Interestingly, there was a trend towards a decrease in myelination in Df1 cells, which had a smaller proportion of myelinated segments (**Figure 5b, g-j**). Branch order myelination did not show any significant alteration and PV^+^ interneurons were proportionally more myelinated in higher branch orders (**Figure 5c**), as previously described in PV^+^ cells^44^. The total myelin length in distal branch order axonal segments trended towards a decrease in Df1 PV^+^ interneurons, although the percentage of myelinated axon was not significantly different (**Figure 5e, f**), suggesting that this trending increment of myelination is due to longer axonal segments in control cells.

**Figure 5.**
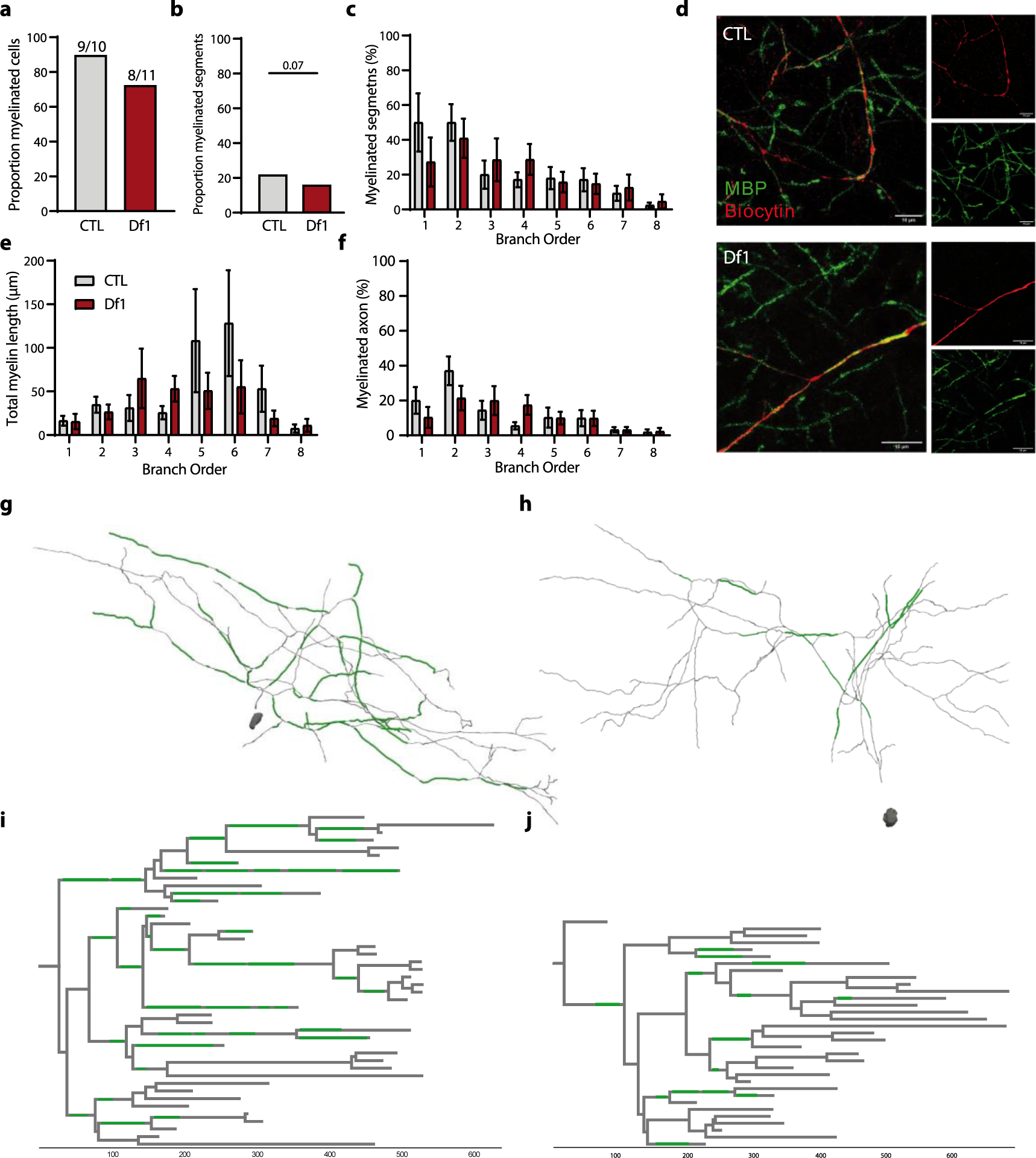
PV^+^ interneurons in Df1 mice are myelinated. **a)** Proportion of myelinated PV^+^ interneurons in Df1 and control littermates (Fisher’s exact test p = 0.59) **b)** The proportion of myelinated segments in each condition (Fisher’s exact test p = 0.078). **c)** Percentage of myelinated segments per branch order (Two-way ANOVA; F (7, 137) = 0.69; p = 0.67). **d)** Image at high magnification of a control and Df1 myelinated trace (axon in red; myelin in green). **e)** Total myelin length per branch order (Two-way ANOVA; F (7,146) = 1.05; p = 1.05). **f)** Percentage of myelinated axon per branch order (Two-way ANOVA; F (7, 138) = 1.26; p = 0.39). **g, h)** Reconstruction of a control (left) and a Df1 PV+ (right) cell’s axon with myelinated segments (green). **i, j)** Axonograms of the corresponding cells above.

## Discussion

In this study, we employed a mouse model of the 22q11DS to better understand the underlying pathophysiology in schizophrenia in different brain areas. We found increased hyperexcitability of PV^+^ interneurons in the dCA1 region of the hippocampus, while PV^+^ interneurons in the PFC were not affected. These alterations in membrane properties were masked when applying synaptic blockers, indicating that synaptic transmission is involved in the observed changes in excitability of PV neurons. Furthermore, PV^+^ interneurons showed a decrease in the expression and functionality of a subset of potassium channels in this region, the Kv1.1 channel, known for its role in controlling PV^+^ interneuron excitability but also for its involvement in synaptic transmission^29,45^. Activation of the channel with a specific agonist restored the excitability levels of PV^+^ cells. Lastly, myelination was not a significantly altered, suggesting that other intrinsic elements or extrinsic inputs are more involved in the changes observed in the Df1 mouse model.

### Hyperexcitability of PV^+^ interneurons: significance and causes

PV^+^ interneurons have been widely studied in schizophrenia. Different alterations have been described in literature corresponding to decreased levels of GABAergic mRNAs^46^ and hypofunction of these cells^16,23,25^, leading to an overall hyperactivity of brain networks. The hippocampus has always been central to the neuropathology of schizophrenia. Previous research had established that this region is hyperactive in psychotic syndromes^47^, corresponding to a dysregulation of glutamate neurotransmission or a hypofunction of the inhibitory system^23,24,47^. Conversely, we found that the electrophysiological properties of PV^+^ interneurons led to a hyperexcitability of these cells, where rheobase and AP threshold were decreased and firing frequency was correspondingly increased. Implications on the general excitability of the hippocampus were not further studied, although hypothetically an increment of the excitability of PV^+^ cells could promote a decrease of the overall excitability. Numerous reasons can lie behind this differential alteration.

(1) The areas of the hippocampal structure present different cellular composition and activity depending on the functions and the role they play. The CA1 region is highly involved in the formation, consolidation and retrieval of memories^48^. Our study focuses on the dCA1, involved in primary cognitive functions and spatial memory, whilst the majority have mainly looked at the vCA1, which exerts a role in emotions^49^. Other studies suggested that the enrichment of encoded excitatory genes in CA3 region might be a big contributor for the hippocampal hyperactivity^50^, instead of the general PV^+^ interneuron population being the major cause. Interestingly, many studies report reduction in the number of PV^+^ interneurons in CA2 but not in CA1 region^24,26^, in line with our observations. Further studies investigating whether PV^+^ interneurons from different subregions are affected differently in schizophrenia would hence be of great relevance for the field. (2) Mice harbouring the 22q11 mutation have helped understanding the genetic contribution of the pool of genes to the development of the disease. Different mouse models have been created with the aim to study 22q11DS. These models comprise longer or shorter regions of the deletion locus^51^, which may affect different genes that entails distinct phenotypes. Therefore, the analysis of individual models is important to understand the different mutations and phenotypes that this deletion encodes. Whether our findings are specific to these model or other models might show similar impairments, requires of more exhaustive analysis of the genes encoded in the Df1 region. (3) Schizophrenia is linked to alterations in the maturation of GABAergic neurons in the PFC during adolescence^24,52,53^. However, hippocampal PV^+^ maturation has also been put forward as crucial for cognitive functions and hippocampal-dependent memory ^54^. Our study shows that hippocampal PV^+^ neurons from young adult Df1 mice present aberrant excitability. We propose that PV^+^ interneuron exhibit a transient increase in their excitability, due to alterations in their channel properties, that might be followed by a chronic hypoexcitability and an overall hyperactive hippocampal network, as reported in various studies^25^. Whether the hyperexcitability can be reversed at an earlier time-point and which are the implications in the overall connectivity of the brain are out of the scope of this study and remains to be explored. (4) Synaptic inputs had an effect on the excitability of PV^+^ interneurons in Df1mice. After bath application of synaptic blockers, the increment of PV^+^ excitability was lost, indicating that synaptic inputs play a role in the modulation of excitability of these cells. Similarly, in *Lgdel/+* mice, the impairment in PV^+^ firing activity was also reversed when excitatory and inhibitory inputs were blocked^23^. Nonetheless, we did not observe changes in sIPSC or sEPSC, in opposition to what was reported in *Lgdel/+* mice^23^. Intrinsic excitability is widely controlled by synaptic transmission and both EPSC and IPSC inputs have been shown to regulate neuronal excitability in various ways^55^, suggesting that synaptic plasticity might be modulating the alterations observed in GABAergic excitability.

### Potassium channel defects in 22q11DS

The organisation of voltage-gated potassium channels (K_v_) along the AIS of interneurons allows for the control of AP properties and signalling^27^. K_v_1.1 channels are not only located in the AIS but also in the presynaptic terminals of the fast-spiking interneurons^37^. Despite not finding alterations in AIS morphology, K_v_1.1 distribution along the AIS could be a better estimate of potassium abnormalities in PV^+^ interneurons. K_v_1.1channels are involved in the regulation of neuronal excitability. Previous studies have demonstrated that blocking K_v_1.1 channels in hippocampal circuits leads to increased firing frequency, decreased rheobase and AP threshold and increased half-width of PV neurons^29,37^, consistent with our results. Moreover, we observed a substantial decrease of somatic K_v_1.1 channel expression in PV^+^ cells in the hippocampus, accompanied by a reduced Kv1.1-specific potassium current near the activation threshold of these channels. Intriguingly, activation of K_v_1.1 channels with 4TFMP had divergent results when the drug was applied alongside synaptic blockers or without. Under baseline conditions, we observed a rescue of PV^+^ interneurons excitability upon application of 4TFMP in Df1 mice, while the same conditions yielded no changes in PV^+^ neurons from control mice. On the other hand, when synaptic activity was blocked, 4TFMP had an effect on PV^+^ interneuron’s excitability in control mice but not in Df1 mice. This results, along with the findings in Supplementary figure 2, expose a differential response of PV^+^ neuron membrane properties to synaptic blockers in control versus Df1 mice, and highlight a clear involvement of presynaptic K_v_1.1 channels in our observed phenotype. Zbili and colleagues previously demonstrated that while a reduction in K_v_1.1 channel density at the axon initial segment can contribute to enhanced PV^+^ interneuron excitability, a loss of presynaptic K_v_1.1 function can in turn lead to increased synaptic transmission in the hippocampus^37^. Future studies investigating how these two processes are involved in the overall dysfunction of PV^+^ interneurons in SCZ model would be of great relevance.

Furthermore, other potassium channels, such as K_v_3.1 are also expressed in the soma and axon terminals of PV^+^ interneurons and are of great importance for the firing frequency of these cells^31^. Interestingly, reductions in this channel is linked to schizophrenia patients^56^. Given their role in both intrinsic and synaptic homeostatic plasticity of PV neurons, K_v_3 channels could also be underlying PV neuron dysfunction in our model^57^. Future investigation could tackle the involvement of this particular channel in the excitability of PV^+^ interneurons in schizophrenia.

### Myelination of PV+ interneurons in schizophrenia

The myelin hypothesis in schizophrenia has emerged in the last few years, when they observed that patients presented a reduction of white matter^58,59^.Individual genes have been studied in mouse models to facilitate the understanding of each candidate on the disease. One of the most examined genes in this cluster is Tbx1. This gene is a transcription factor responsible for the regulation expression of thousands of (congenital) genes^60^. Interestingly, Tbx1 has been linked to alterations in myelination, where they found a reduction in the number of large myelinated axons but increased g-ratios^61^, proposing a correlation between 22q11DS genes and myelination alterations. Notably, PV^+^ interneurons are highly myelinated in the cortex and hippocampus^43,44^. The relationship between PV^+^ dysfunctionality in schizophrenia and myelination^17^ attracts to study whether 22q11DS mouse models present also alterations in the myelination profile of these cells. We identified that the proportion of myelinated PV cells was not changed. However, there was a trend towards a decrease in the percentage of myelinated segments in Df1 cells. This trend was only observed in the proximal axon, where most of the myelination occurs^44^. Whether myelination deficits may follow at a later stage or as a consequence of the sustained chronic hyperexcitability of PV^+^ interneurons, or whether pyramidal neurons are in fact affected, should be addressed in future studies.

Together, this study exposes an increase in PV^+^ neuronal excitability in the hippocampus of mice harbouring the 22q11DS. This excitability has a synaptic component that can be derived, among others, by K_v_1.1 channel deficiencies and synaptic transmission. Activation of these potassium channels can partially recover the intrinsic properties, and opens new targets for therapeutic studies.

## Material and methods

### Mice

All experiments were conducted under the approval of the Dutch Ethical Committee and in accordance with the Institutional Animal Care and Use Committee (IACUC) guidelines.

For this study, we obtained two mouse lines from Jackson Laboratory: B6J-Del(16Es2el-Ufd1l)217Bld (referred as Df1). This line was crossed with a Tg(Pvalb-tdTomato)15Gfng/J (referred as PVtomato) line in order to visualize a specific subset of neurons. The resulting line was B6.Del(16Es2el-Ufd1l)217Bld x -Tg(Pvalb-tdTomato)15Gfng/J (referred as Df1::PVtom). Mice employed were heterozygous for the deletion, just as human 22q11DS patients. All mutant lines were viable and healthy. No behavioural abnormalities and no spontaneous seizures were ever observed.

Mice between 7 and 14 weeks old from both sexes were used in these experiments. Animals were group-housed and maintained on a regular 12 h light/dark cycle at 22 °C (±2 °C) with access to food and water *ad libitum*.

### Electrophysiology

Mice were anaesthetised under 5% isofluorane and decapitated. Brains were removed in ice-cold partial sucrose-based solution containing (in mM): sucrose 70, NaCl 70, NaHCO_3_ 25, KCl 2.5, NaH_2_PO_4_ 1.25, CaCl_2_ 1, MgSO_4_ 5, sodium ascorbate 1, sodium pyruvate 3 and D(+)-glucose 25 (carboxygenated with 5% CO_2_/95% O_2_). Coronal slices of 300 µm thick from the hippocampus were obtained with a vibrating slicer (Microm VT1200 S, Leica) and incubated for 45 min at 37 °C in holding artificial cerebro-spinal fluid (ACSF) containing (in mM): 127 NaCl, 25 NaHCO_3_, 25 D(+)-glucose, 2.5 KCl, 1.25 NaH_2_PO_4_, 1.5 MgSO_4_, 1.6 CaCl_2_, 3 sodium pyruvate, 1 sodium ascorbate and 1 MgCl_2_ (carboxygenated with 5% CO_2_/95% O_2_). Slices were then transferred into the recording chamber where they were continuously perfused with recording ACSF (in mM): 127 NaCl, 25 NaHCO_3_, 25 D-glucose, 2.5 KCl, 1.25 NaH_2_PO_4_, 1.5 MgSO_4_ and 1.6 CaCl_2_. For the synaptic blocker experiments, a cocktail containing 50 µM AP-5 (HelloBio), 10 µM CNQX (HelloBio) and 10 µM Gabazine (HelloBio) was added into the recording ACSF. For the activator experiments 100 µM 4TFMPG (Thermo Fisher) was added into the recording ACSF solution and cells were perfused with this solution for 10 minutes and then washed with normal recording solution.

Cells were visualized using an upright microscope (BX51WI, Olympus Nederland) equipped with oblique illumination optics (WI-OBCD; numerical aperture 0.8) and a 40x water-immersion objective. Images were collected by a CCD camera (CoolSMAP EZ, Photometrics) regulated by Prairie View Imaging software (Bruker). PV^+^ interneurons were visualised using a DsRed filter (Semrock, Rochester, NY, USA). Electrophysiological recordings were acquired using HEKA EPC10 quattro amplifiers and Patchmaster software (40 Hz sampling rate) at 33°C. Patch pipettes were pulled from borosilicate glass (Harvard Apparatus) with an open tip of 3-5.5 MΩ of resistance and filled with intracellular solution containing (in mM) 125 K-gluconate, 10 NaCl, 2 Mg-ATP, 0.2 EGTA, 0.3 Na-GTP, 10 HEPES and 10 K2-phosphocreatine, pH 7.4, adjusted with KOH (280 mOsmol/kg), with 5 mg/mL biocytin to fill the cells. Series resistance was kept under 20 MΩ with correct bridge balance and capacitance fully compensated; cells that exceeded this value were excluded from the study. sEPSC were recorded at -70 mV holding potential during 5 min. For sIPSC and autaptic neurotransmission recordings, patch pipettes were filled with a high [Cl−] intracellular solution containing (in mM) 70 K-gluconate, 70 KCl, 10 HEPES, 1 EGTA, 2 MgCl2, 2 MgATP, and 0.4 NaGTP; pH 7.2 adjusted with KOH; 285 mOsm. The estimated ECl was around –16 mV based on the Nernst equation. AMPA- and NMDA-mediated currents were excluded using DNQX (10 μM) and D-AP5 (50 uM), respectively in the bath perfusion. Gabazine (10 μM) was also applied by bath perfusion to block GABA_A_Rs. Neurons were kept at -70 mV, and unstable cells were not considered in the analysis. Cells were filled with biocytin for at least 20 min. For autaptic responses, PV+ interneurons were clamped at -80 mV. Brief depolarizing voltage steps (0.2-0.6 ms, from -80mV to 0 mV) were injected in to induce a GABAergic response that was abolished by gabazine

Data analysis was performed offline using Igor Pro 9.00 with a custom-designed script. Passive and active properties were derived from voltage responses to current injection pulses of 500 ms duration in 100 pA intervals ranging from -200 to +750 pA. The AP waveform and physiological characteristic were determined from the first elicited AP. Synaptic events were detected using Mini Analysis Program (Synaptosoft).

### Voltage clamp

Slices were perfused using recording ACSF including 1 µM tetradotoxin (TTX; HelloBio). Cells were then perfused with a solution containing 100 nM of the K_v_1.1 specific blocker DTx-K (Alomone labs) and washed for 6-8 minutes. Recordings were performed 4 minutes after the solution had entered the bath, for maximum efficacy of the blocker. Voltage clamp analysis was performed using Heka online analysis. Leak currents were automatically detected and substracted online from the analysis. DTx-K currents were substracted from the general potassium currents to asses K_v_1.1-specific potassium currents.

### Immunohistochemistry

Df1 mice and their control littermates were anaesthetized intraperitoneally with pentobarbital natrium (Nembutol®) before performance of cardial perfusion using 4 % formaldehyde (PFA). Brains were removed and fixed in 4% PFA for 2 h at room temperature. Next, brains were transferred into 10% sucrose phosphate buffer (0.1 M PB) and stored overnight at 4°C. To improve the slicing process, brains were embedded in 12% gelatin-10% sucrose blocks and left during 1.5 h at room temperature in 10% paraformaldehyde-30% sucrose solution (PB 0.1 M) and later incubated overnight in 30% sucrose solution at 4°C. Brains were then sliced in coronal slices at 40 µm thick using a freezing microtome (Leica, Wetzlar, Germany; SM 2000R) and stored in 0.1 M PB.

For the staining, slices were blocked with 0.5% Triton X-100% (MerkMillipore) and 10% normal horse serum (NHS; Invitrogen, Bleiswijk, The Netherlands) for 1 h at room temperature, and incubated over 72 h at 4°C with primary antibodies mouse anti-K_v_1.1 (1:300, NeuroMab clone K36/15), rabbit anti-PV (1:1000, Swant PV25) and rabbit anti-AnkG (1:300, Synaptic Systems) in PBS buffer containing 0.4% Triton X-100 and 2% NHS. Secondary antibodies anti-mouse Alexa 488 (1:300, Invitrogen), anti-rabbit Cy3 (1:300, Invitrogen) and anti-rabbit Alexa 647 (1:300, Invitrogen) were employed for 2 h at room temperature. Slices were coverslipped with Vectashield H1000 fluorescent mounting medium (Vector Labs, Peterborough, UK).

### Mice biotin-filled cells

Biocytin-filled cells were fixed in 4% PFA overnight and then stored in PBS-Na azide 2%. Slices were then stained with streptavidin-Cy3 secondary antibody (1:300; Invitrogen) in 0.4% Triton X-100 and 2% NHS PBS solution during 3 h and then coverslipped with Mowiol (Sigma). After confocal imaging, cells were unsealed and rinsed in 30% sucrose before resectioning. Slices were resectioned in 40 µm thick slices and then stained with primary antibody mouse anti-MBP mouse anti-MBP (1:300, Santa Cruz, F-6, sc-271524) for 72 h at 4°C and secondary antibody anti-mouse Alexa-488 (1:300, Invitrogen).

### Western Blot

Mice were anaesthetised using isufluorane and hippocampal tissue was isolated from the rest of the brain. Samples were sonicated in lysis buffer (0.1 M Tris-HCl [pH 6.8], 4% SDS) and protease inhibitors (P8340, Sigma). We determined the protein concentration of the supernatant using a BCA protein assay (Pierce). 10 µg of the sample were loaded into the running gel (3450124, Bio-rad), that was later transferred onto a nitrocellulose membrane (1704159, Bio-rad). Blots were incubated with primary antibodies anti-K_v_1.1 (NeuroMab, clone K36/15) and anti-K_v_3.1 (NeuroMab clone N16b/8) with a concentration of 1:1000 overnight at 4°C. Next, blots were incubated with secondary antibody GFP goat anti-mouse (Licor, 1:10000) for 1 h at room temperature. To normalise protein levels, we used primary antibody anti-actin (). Blots were imaged using using an Odyssey® Imager (Licor) and analysed offline with Image Studio™ Lite (Licor).

### Confocal imaging and reconstruction

Images were taken using a Zeiss LSM 700 microscope (Carl Zeiss) equipped with Plan-Apochromat 10x/0.45 NA, 40x/1.3 NA (oil immersion) and 63x/1.4 NA (oil immersion) objectives. Secondary fluorophores were imaged using excitation wavelengths of 488 nm, 555 nm, and 639 nm. A whole-cell overview image of the biocytin-filled cells was taken with the 63x objective for reconstruction purposes. After resection and staining with MBP, slices were imaged with a 40x magnification.

Whole-cell overview images were then transferred into Neurolucida 360 software (v2.8; MBF Bioscience). The axon was reconstructed using the interactive tracing with the Directional Kernels method and myelinated segments were added post-hoc. Axons were considered as myelinated when they presented at least one MBP-positive internode and unmyelinated when no MBP-positive internode could be identified up to at least 9^th^ branch order. Analysis and quantification were done using Neurolucida Explorer (MBF Bioscience). No spatial corrections were made for tissue shrinkage.

### Confocal imaging and fluorescence analysis

Confocal pictures of PV, Kv1.1 and AnkG stainings were taken using objectives of 20x/0.45 NA and 40x/1.3 NA (oil immersion). 20x pictures were used for quantification of cell numbers, and 40x z-stack confocal picture were used for fluorescence analysis. Three stacks were quantified per mouse and hemisphere. Selected cells were manually outlined and fluorescence of individual cells was measured using the measure stack plugin in Fiji (ImageJ 5.12h). Soma was removed and used as a background noise.

### Statistical analysis

All statistical analysis were operated using GraphPad Prism 8. Data was firstly analysed for normality using X. No outlier was identified or removed. Data sets following normal distribution were analysed using unpaired two-tailed *t*-test or a two-way ANOVA and the corresponding post hoc test. Data sets without a normal distribution were analysed using Mann Whitney test. Graphs were also made using GraphPad Prism 8.

## Supplementary data

**Supplementary figure 2.**
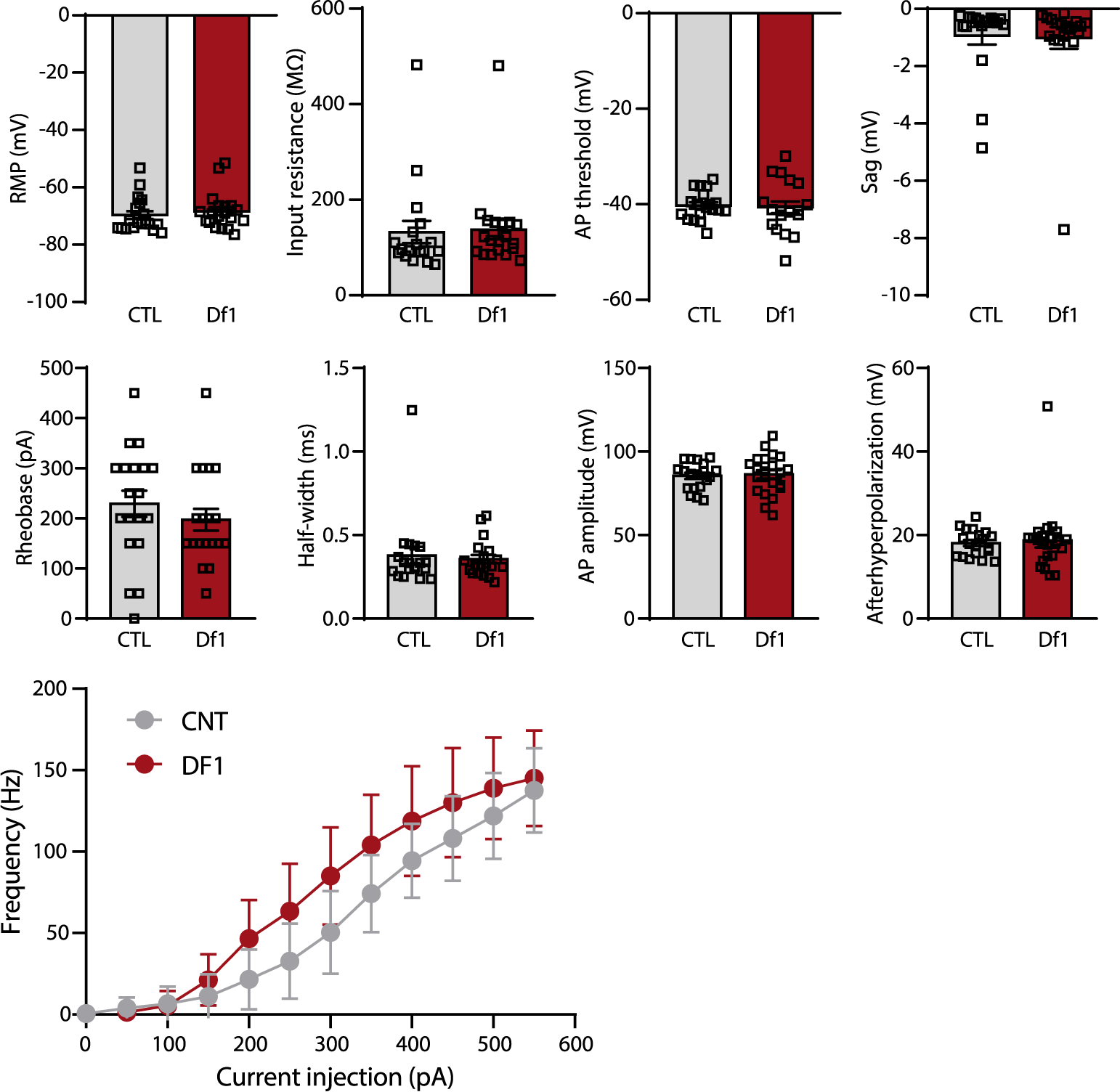
Increased excitability of prefrontal PV^+^ interneurons in Df1 mice. **a)** Resting membrane potential (t-test: t = 0.65; p-value = 0.52; CTL average: -69.66 mV; Df1 average: - 68.34). **b)** Input resistance (t-test: t = 0.16; p-value = 0.87; CTL average: 132.5 MΩ; Df1 average: 137.4 MΩ). **c)** AP threshold (t-test: t = 0.28; p-value = 0.78; CTL average: -40.25 mV; Df1 average: -40.65 mV). **d)** Sag (t-test: t = 0.87; p-value = 0.16; CTL average: -0.93 mV; Df1 average: -1.09 mV). **e)** Rheobase (t-test: t = 0.92; p-value = 0.36; CTL average: 228.9 pA; Df1 average: 197.4 pA). **f)** Half-width (Mann-Whitney: U = 176; p-value = 0.90; CTL average: 0.38 ms; Df1 average: 0.36 ms). **g)** The amplitude of the AP (t-test: t = 0.27; p-value = 0.78; CTL average: 85.39 mV; Df1 average: 86.27 mV). **h)** The afterhyperpolarisation (Mann-Whitney: U = 198; p-value = 0.79; CTL average: 18.08 mV; Df1 average: 18.74 mV). **i)** Firing frequency in response to depolarising steps from 0 to 550 mV during 500 ms.

**Supplementary figure 2.**
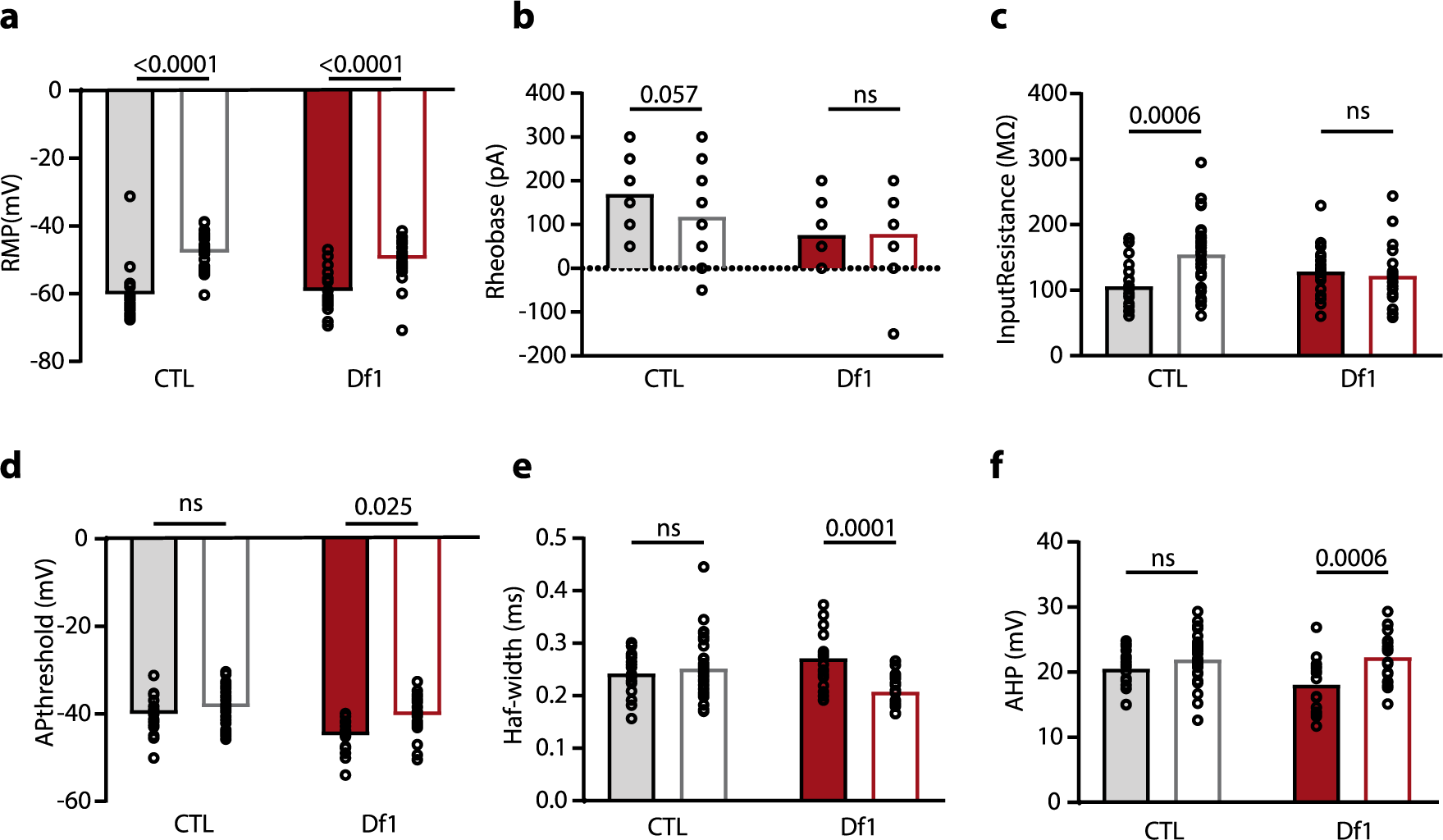
Comparison between intrinsic properties of PV^+^ interneurons with and without blockers. **a)** RMP was significantly increased in both groups after application of synaptic blockers (Two-way ANOVA: F (1,90) = 1.176; p-value = 0.28; Šidák’s multiple comparisons test; CNT, p < 0.0001; Df1, p < 0.0001) **b)** Rheobase was not affected in the presence of synaptic blockers (Two-way ANOVA: F (1, 86) = 2.43; p-value = 0.12; Šidák’s multiple comparisons test; CNT, p = 0.057; Df1, p = 0.99). **c)** Input resistance was increased in the presence of synaptic blockers in CTL cells (Two-way ANOVA: F (1, 87) = 8.21; p-value = 0.005; Šidák’s multiple comparisons test; CNT, p = 0.0006; Df1, p = 0.87) **d)** AP threshold was significantly affected by synaptic blockers in Df1 group (Two-way ANOVA: F (1, 86) = 2.6; p-value = 0.11; Šidák’s multiple comparisons test; CNT, p = 0.40; Df1, p = 0.0025). **e)** HW was significantly decreased by synaptic blockers in Df1 group (Two-way ANOVA: F (1, 87) = 12.83; p-value = 0.0006; Šidák’s multiple comparisons test; CNT, p = 0.75; Df1, p = 0.0001). **f)** AHP was significantly increased by synaptic blockers in Df1 group (Two-way ANOVA: F (1, 87) = 3.48; p-value = 0.065; Šidák’s multiple comparisons test; CNT, p = 0.34; Df1, p = 0.0006).

**Supplementary figure 3.**
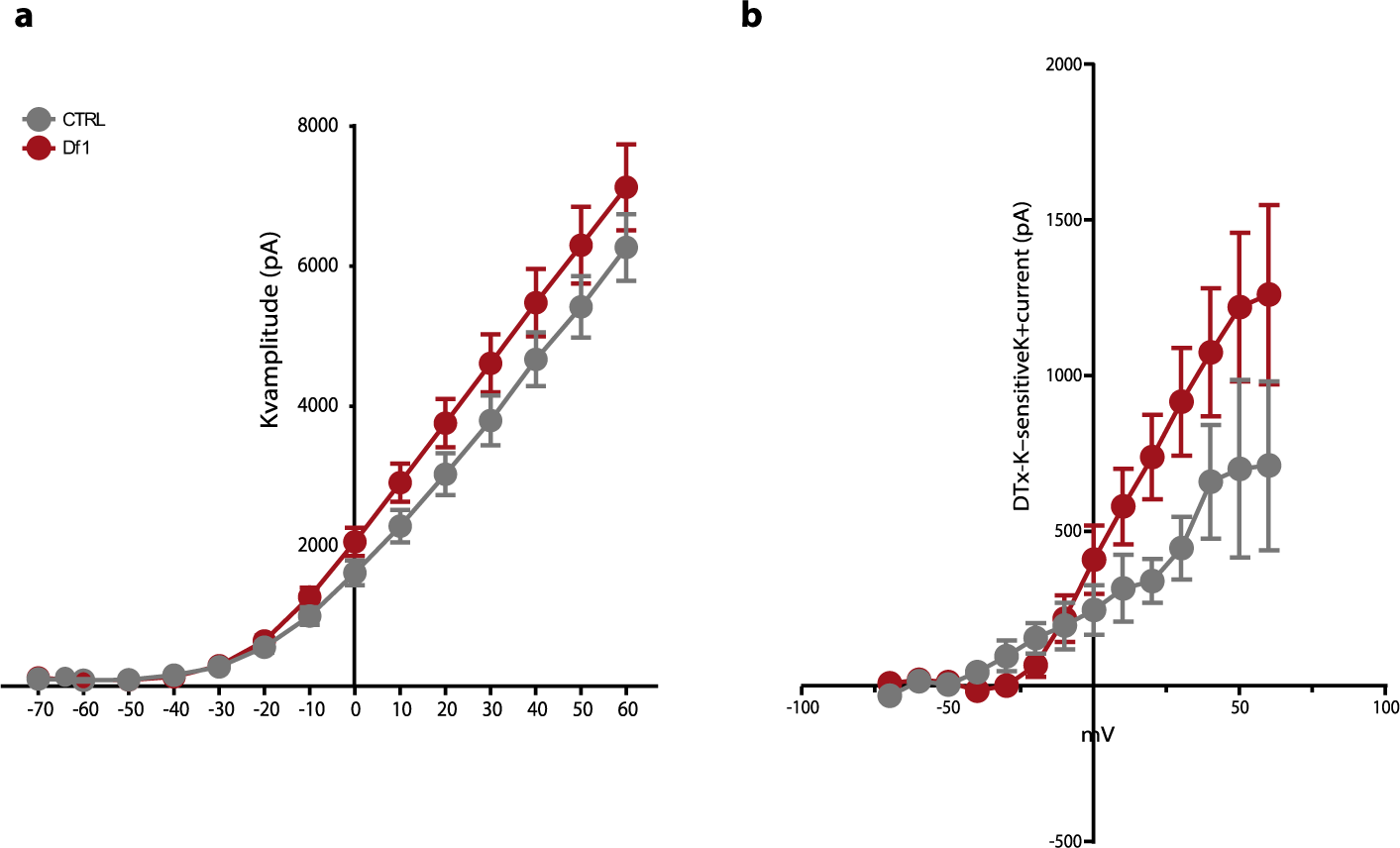
Potassium current plots. **a)** Plot of the voltage dependence of the potassium-sensitive currents in Df1 and control PV+ interneurons (ANOVA: F (13, 338) = 1.73; p-value = 0.052). **b)** Plot of the voltage dependence of the DTx-K-sensitive currents in Df1 and control PV+ interneurons (ANOVA: F (13, 169) = 2.35; p-value = 0.006).

**Supplementary figure 4.**
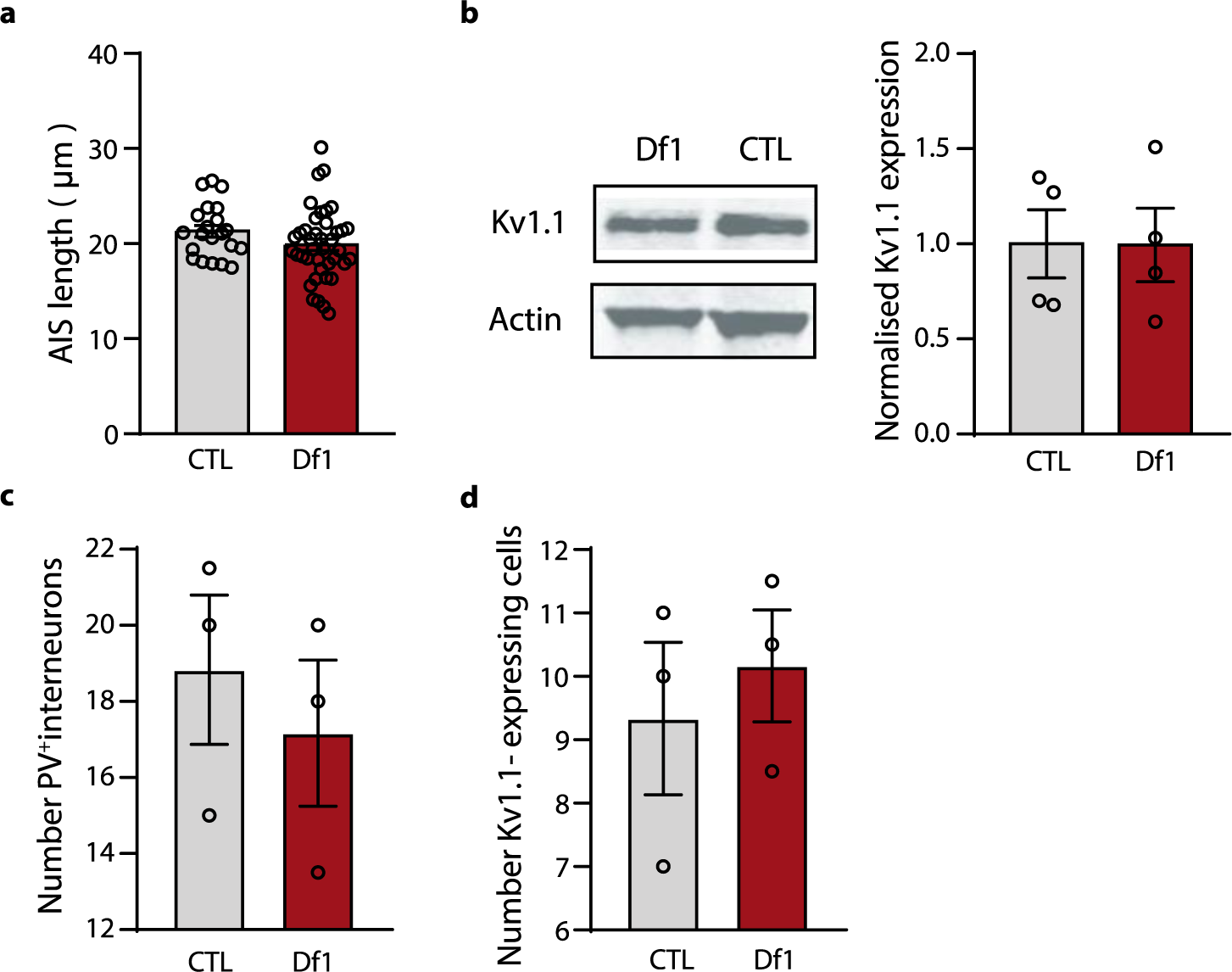
Molecular characteristics of Df1 hippocampal cells. **a)** Axon initial segment length in PV+ interneurons in dCA1 region (t-test: t = 1.52; p = 0.13). **b)** Western blot expression of Kv1.1 channels in the hippocampus (t-test: t = 1.02; p = 0.98). **c)** Number of PV^+^ interneurons in the dCA1 (t-test: t = 1.04; p = 0.6). **d)** Number of Kv1.1-expressing cells in the dCA1 (t-test: t = 1.86; p = 0.6).

**Supplementary figure 5.**
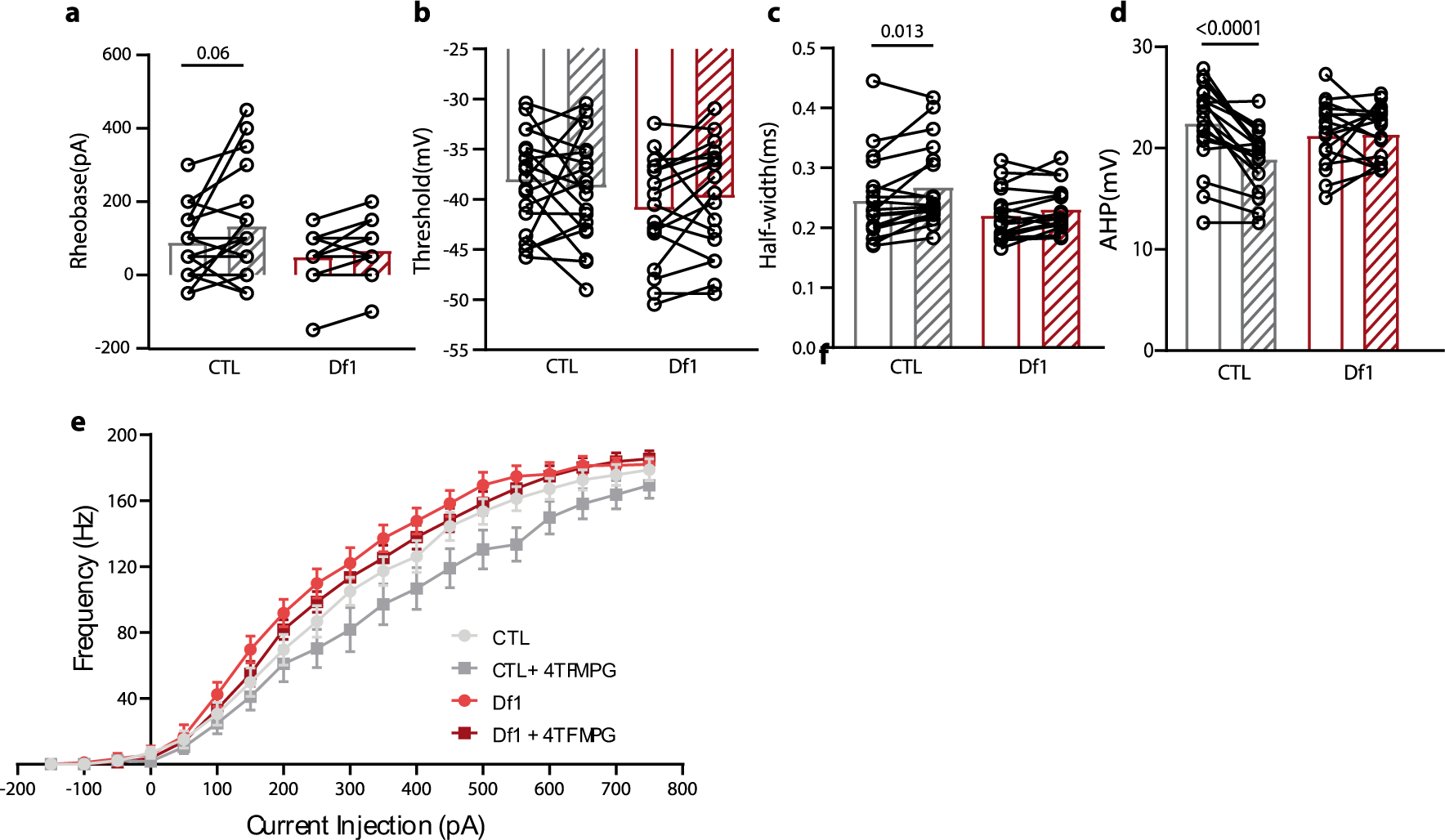
4TFMPG selective agonist for Kv1.1 does not have an effect on PV^+^ interneurons in the presence of synaptic blockers. **a)** Rheobase was not significantly changed in either group after application of 4TFMPG (Two-way ANOVA: F (1, 34) = 0.86; p-value = 0.36; Šidák’s multiple comparisons test; CNT, p = 0.06; Df1, p = 0.65) **b)** AP threshold was not affected after application of 4TFMPG (Two-way ANOVA: F (1, 34) = 1.35; p-value = 0.25; Šidák’s multiple comparisons test; CNT, p = 0.85; Df1, p = 0.47). **c)** Half width was increased in the control group bath application of 4TFMPG (Two-way ANOVA: F (1, 34) = 1.88; p-value = 0.18; Šidák’s multiple comparisons test; CNT, p = 0.013; Df1, p = 0.20) **d)** AHP was reduced only in control cells in the presence of 4TFMPG (Two-way ANOVA: F (1, 34) = 14.17; p-value = 0.0006; Šidák’s multiple comparisons test; CNT, p < 0.0001; Df1, p = 0.98) **e)** Firing frequency in response to depolarising steps from 0 to 750 mV during 750 ms before and after bath application of the 4TFMPG (Two-way ANOVA: F (54, 1287) = 18.09; p-value < 0.0001).

**Supplementary figure 6.**
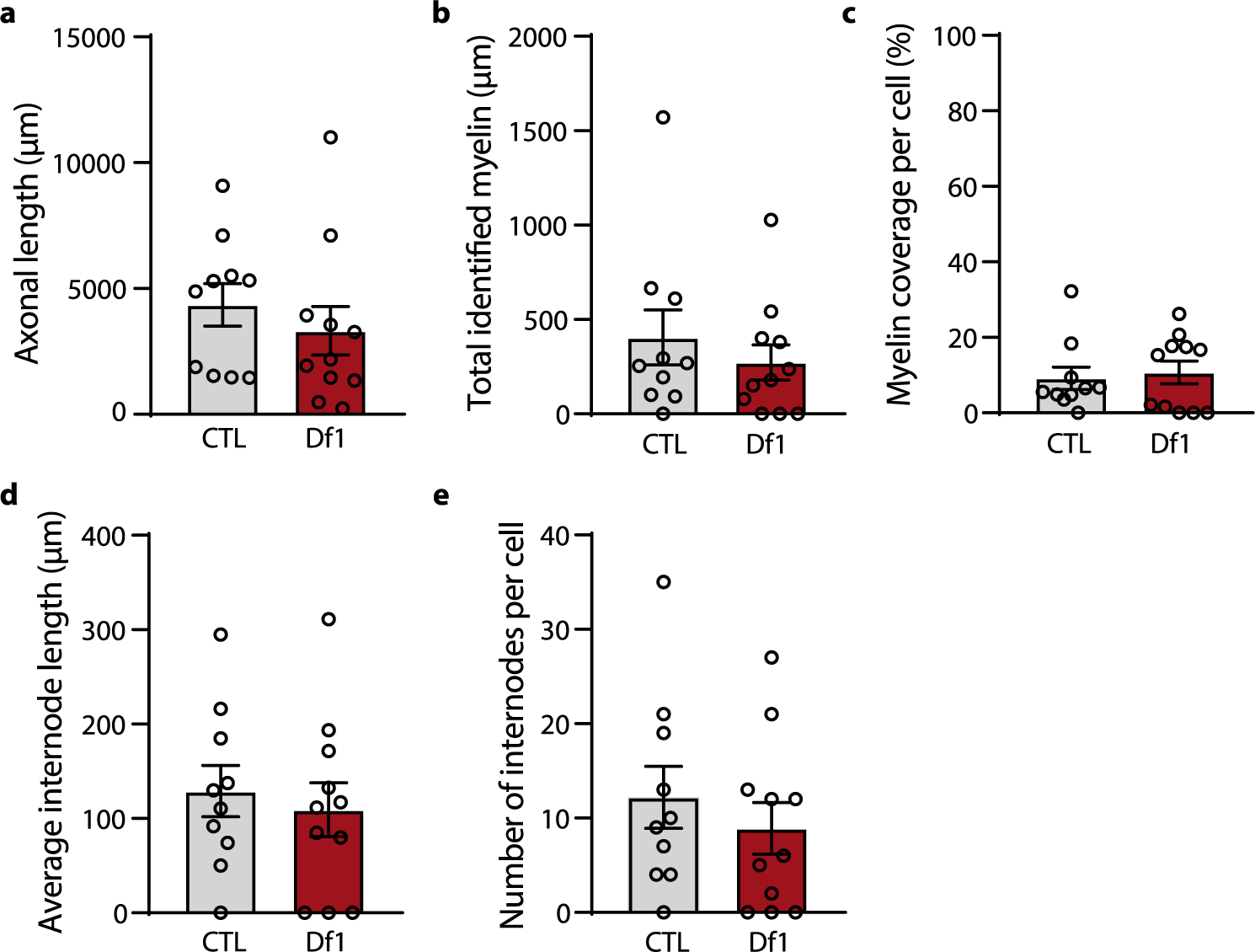
Axonal and myelin analysis of PV+ interneurons. **a)** Axonal length (t-test: t = 0.8; p = 0.43). **b)** Total amount of myelin (t-test: t = 0.78; p = 0.45). **c)** Percentage of myelination per cell (t-test: t = 0.36; p = 0.72). **d)** Average internode length (t-test: t = 0.49; p = 0.62). **e)** Number of internodes (t-test: t = 0.77; p = 0.45).

